# Expression of normal levels of the Pumilio domain protein PUF3 is required for optimal growth of *Trypanosoma brucei*

**DOI:** 10.1101/682542

**Authors:** Kevin Kamanyi Marucha, Christine Clayton

## Abstract

The *Trypanosoma brucei* pumilio domain protein PUF3 is a cytosolic mRNA-binding protein that suppresses expression when tethered to a reporter mRNA. An induced reduction of PUF3 in bloodstream forms caused a slight growth defect and slightly delayed differentiation to the procyclic form, but the cells lost both defects upon prolonged cultivation. Both *PUF3* genes could also be deleted in bloodstream-form and procyclic-form trypanosomes, suggesting that *in vitro*, at least, these life-cycle stages do not require PUF3. Procyclic forms without PUF3 grew somewhat slower than wild-type, but were able to transform to bloodstream forms after induced expression of the bloodstream-form RNA-binding protein RBP10. In contrast, ectopic expression of C-terminally tagged PUF3 in procyclic forms impaired viability. There was little evidence for specific binding of PUF3 to bloodstream-form mRNAs and RNAi had no significant effect on the transcriptome. Moreover, mass spectrometry revealed no PUF3 binding partners that might explain its suppressive activity. Since PUF3 is conserved in all Kinetoplastids, we suggest that it might be required within the invertebrate host, or perhaps implicated in fine-tuning gene expression.

## Introduction

Pumilio domain proteins contain multiple “Puf” repeats, each of which can bind to a single nucleotide in RNA. Classically, there are 8 adjoined repeats, which together bind to a sequence with consensus of UGUR [1]. *Trypanosoma brucei* is a unicellular eukaryote of the Excavata supergroup. It multiplies as the bloodstream form in man and ruminants, and as the procyclic form in the midgut of tsetse flies. The genome of *T. brucei* encodes eleven PUF proteins with different domain architectures. Only PUF2, 3, 4, and 6 have the canonical 8 pumilio repeats, and all but PUF3 (Tb927.10.310) have low complexity or coiled coil motifs. Most of the PUF proteins are cytosolic; the exceptions are PUF7, PUF8 and PUF10, which are found in the nucleolus [2]. All except PUF11 and PUF8 are conserved in Kinetoplastea, which includes not only parasitic kinetoplastida but also free-living bodonids (S1 Fig A). Conservation between PUF3 homologues is restricted to the eight repeats, a 17-residue region immedietely C-terminal to the last repeat, and two short regions at the N-terminus (S1 Fig B).

We here describe experiments to investigate the function of *T. brucei* PUF3. PUF3 was already known to bind to mRNA in *T. brucei* bloodstream forms [3] and to repress expression when tethered (via a lambdaN peptide) to the 3’-untranslated region of a reporter mRNA [3, 4]. Results of a high-throughput RNAi screen also suggested that reduced PUF3 expression caused a significant loss of fitness in all life-cycle stages tested (bloodstream forms, and early differentiation to the procyclic form). Our results, however, show that PUF3 is not essential in cultured *T. brucei*, although it may have a role during differentiation.

## Results

### PUF3 is cytosolic and rapidly represses expression

To determine the cellular location of PUF3, we used bloodstream-form trypanosomes expressing N-terminally V5-tagged PUF3 from the endogenous locus (V5-PUF3), or trypanosomes with inducible expression of C-terminally myc-tagged PUF3 (PUF3-myc) (Fig 1). By immonofluorescence microscopy, V5-PUF3 was spread throughout the cytosol with no detectable signal from the nucleus (Fig 1A). Cell fractionation using digitonin confirmed that PUF3-myc was not inside mitochondria or glycosomes (Fig 1B). Later, results from high-throughput N-terminal GFP tagging [2] also indicated a cytosolic location.

**Figure 1.**
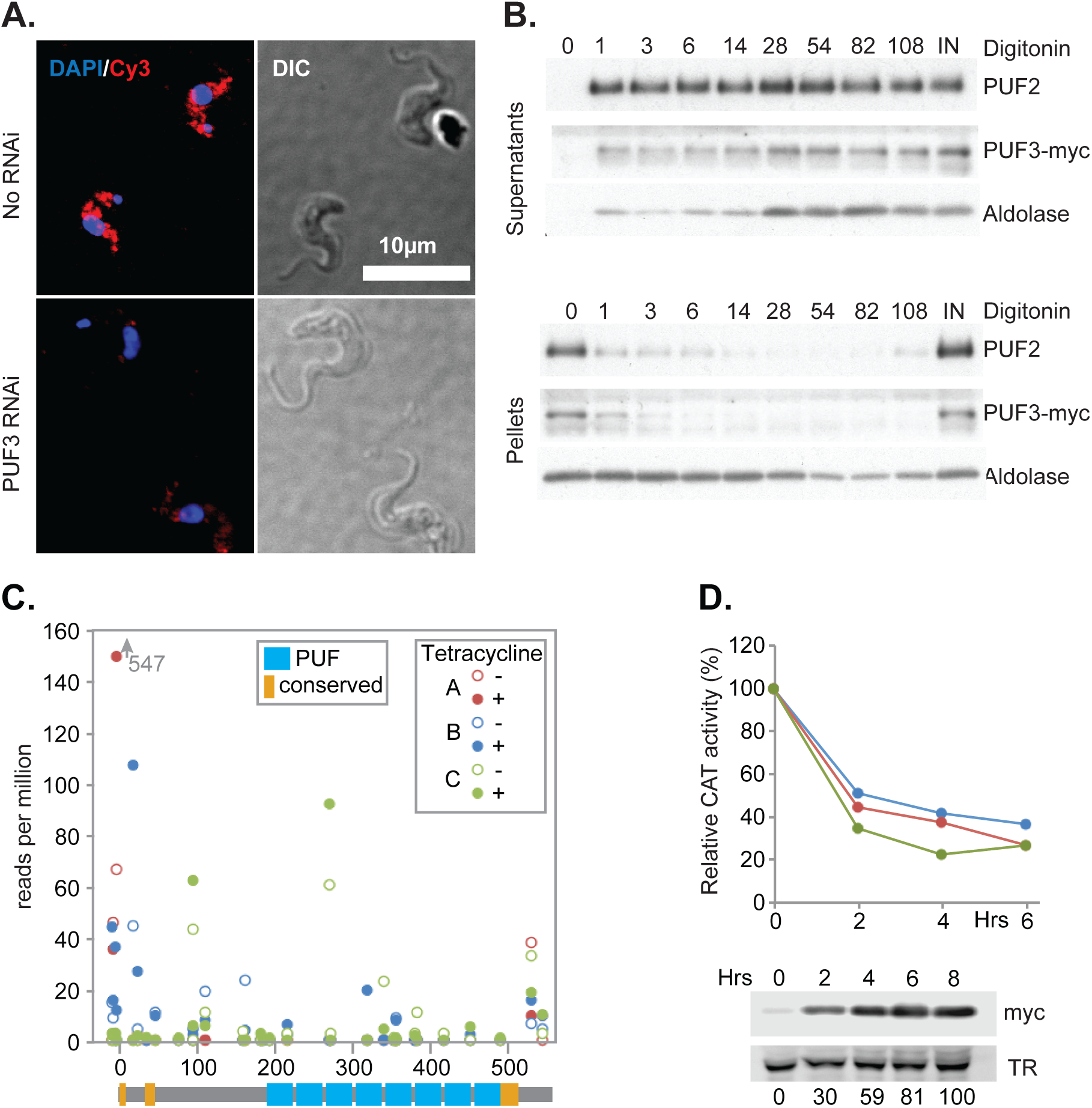
PUF3 location and tethering. **A)** Bloodstream-form trypanosomes with *in situ* tagged PUF3 and inducible RNAi targeting *PUF3* were used. The upper image shows cells without tetracycline and the lower image after 24h tetracycline treatment. Stains are for V5-PUF3 (red) and DNA (blue). DIC is the differential interference contrast. **B)** Cells expressing inducible PUF3-myc (24 h tetracycline) were lysed using increasing concentrations of digitonin. Organelle rupture was detected by the migration of the organelles’ proteins to the supernatant. PUF2 and aldolase were used as cytosolic and glycosomal markers respectively. The amounts of digitonin (mg of digitonin/ mg of protein) are indicated above each lane. **C)** Results of the tethering screen. The bloodstream-form trypanosomes used contained a tetracycline-inducible *PGKB* gene, with five copies of boxB between the open reading frame and the actin 3’-untranslated region (3’-UTR). They were transfected with a library designed for inducible expression of proteins encoded by ∼1kb genomic fragments, preceded by the lambdaN peptide. Tetracycline addition resulted in growth retardation, unless the expressed lambdaN-protein repressed PGKB expression. After induction, cells were grown, then the DNA inserts were amplified and sequenced [4]. This selects for protein sequences capable of suppressing translation and/or inducing mRNA degradation. The graph shows the results for PUF3, with first codon positions on the x-axis, and the reads per million on the Y-axis. Higher values with tetracycline than without tetracycline suggest suppressive activity. The cartoon below shows domains of PUF3, to scale: PUF domains in cyan, and sequences conserved in Kinetoplastea are in orange (S1 Fig B). **D)** Confirmation of tethering results for full-length lambdaN-PUF3-myc. The bloodstream-form trypanosomes used constitutively express a chloramphenicol acetyltransferase (*CAT*) mRNA with five B-boxes between the open reading frame and the actin 3’-UTR. Expression of lambdaN-PUF3-myc was induced by addition of tetracycline and CAT activity was measured. The graph shows results for 3 independent cultures.

Cytosolic phosphoglycerol kinase (PGKB) is toxic to bloodstream-form trypanosomes. Tethering of full-length PUF3 to an mRNA encoding PGKB conferred a selective advantage [3], suggesting that PUF3 suppresses expression. Using random fragments of the *PUF3* gene [4], most selected clones encoded tethered polypeptides that commenced within the first 100 amino acids (Fig 1C). This result suggests that the suppressive sequence is N-terminal to the PUF repeats. We confirmed that tethering of lambdaN-PUF3-myc to a reporter mRNA encoding chloramphenical acetyltransferase (CAT) was repressive: CAT activity decreased within 4 hours (Fig 1D). Consistent with this, PUF3 shows no association with polysomal RNA [5].

### Interactions of TAP-tagged PUF3

To investigate mRNAs bound to PUF3, we made bloodstream-form trypanosomes in which one *PUF3* open reading frame was deleted and the other was fused at the 5’-end with a sequence encoding a tandem affinity purification (TAP) tag. After UV cross-linking, cells were lysed, tagged PUF3 was selected, and bound and unbound RNAs were sequenced for two biological replicates. The unbound RNA fraction was subjected to rRNA depletion using RNase H and oligonucleotides complementary to rRNA [6]. To check the effect of the rRNA procedure on the transcriptome we also sequenced the input RNA from the experiment, either with or without rRNA depletion.

The transcriptome results showed that rRNA depletion had no significant effect on the transcriptome (Fig 2A; Table S1, sheet 3; Fig S2 A). By principal component analysis (Fig S2 A), the unbound RNA from the pull-down experiment clustered with the input samples, which is expected if only a few mRNAs are bound by the selected protein. Rather large numbers of cells were needed to obtain sufficient bound mRNA for standard sequencing library preparation, but these bound fractions were clearly different from the input and unbound RNAs (Fig S2 A). *PUF3* mRNA was enriched in the bound fractions, as expected from affinity purification of the mRNA together with the nascent polypeptide [7],[8],[9]. *7SL* RNA, which is a component of the signal recognition particle, was also, as expected, almost exclusively in the unbound fraction. However, only six mRNAs were more than 2-fold enriched, which could be assigned to chance. The 106 mRNAs that were reproducibly more than 2-fold enriched were not significantly longer than the remaining mRNAs, suggesting that their selection was not due to non-specific RNA binding to PUF3 or the matrix (Fig 2B; Table S1, sheet 2). Long-lived mRNAs were also missing from the enriched fraction (Fig 2C, P=10^−5^), consistent with destabilising activity of PUF3. Since, however, we are not sure whether the tagged PUF3 was fully functional (see below), and enrichment of bound mRNAs was poor, the significance of these results is questionable.

**Figure 2.**
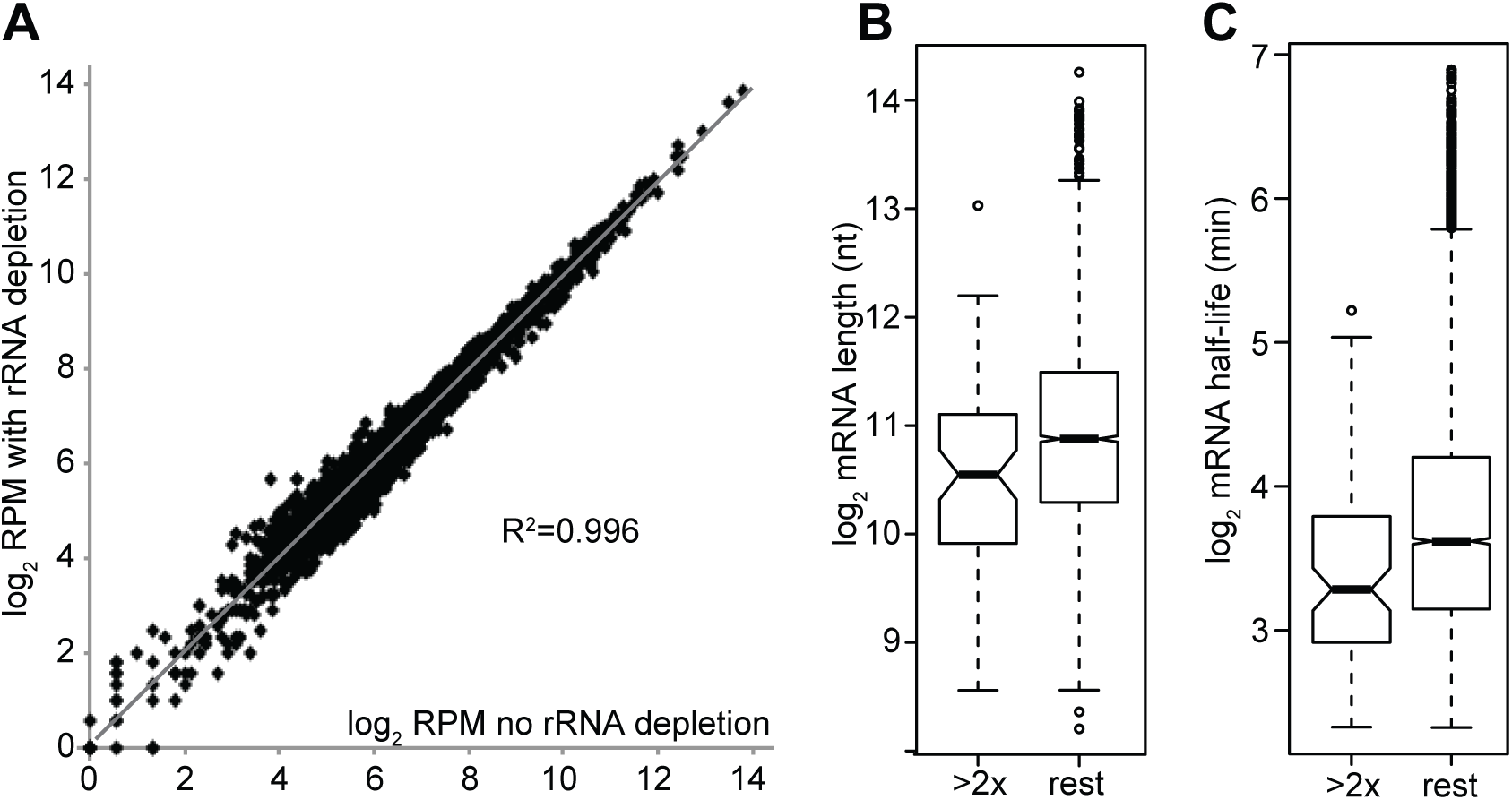
rRNA depletion and RNA pull-down **A**). rRNA depletion using oligonucleotides and RNaseH does not affect the transcriptome. Average reads per million reads for duplicate matched total RNA samples are shown, on a log2 scale. Individual comparisons are in Table S1. **B)** Lengths of bound mRNAs relative to those of the remaining mRNAs. mRNAs were classed as “bound” if both experiments gave a ratio greater than 2. The “bound” mRNAs are not longer than the rest, arguing against non-specific sticking. **C)** Half-lives of bound mRNAs relative to the remaining mRNAs.

We also subjected TAP-PUF3 to tandem affinity purification and identified co-purifying proteins by mass spectrometry (Table S2). The only proteins that appeared to co-purify, relative to TAP-GFP controls, were abundant proteins that had previously been identified in other purifications. The results therefore gave no hint of a mechanism by which PUF3 might inhibit mRNA translation or promote mRNA degradation.

### PUF3 is not required in growing bloodstream forms

To assess the function of PUF3 we first reduced its expression by RNAi. We used two trypanosome cell lines expressing the tet repressor. One, of the EATRO1125 strain, is capable of full differentiation from the bloodstream to the procyclic form. The EATRO1125 bloodstream forms were routinely grown in methyl cellulose-containing medium to maintain differentiation competence. When they reach a density of 2.5-3.0 x 10^6^/ml, they arrest in G_0_, and become “stumpy” - that is, they are shorter and fatter, and express the stumpy-form marker PAD1 and various mitochondrial proteins required for survival in tsetse. When the growth temperature is reduced from 37°C to 27°C and cis-aconitate is added, they differentiate to procyclic forms and start to grow in appropriate medium within 1-2 days. Our Lister 427 strain bloodstream forms, in contrast, can grow to densities of 6-7 x10^6^/ml and never make stumpy forms or PAD1. If these cells are treated at 27°C with cis-aconitate, they are able to express some procyclic-specific mRNAs and proteins, including the procyclic surface protein EP procyclin, but they are unable to grow as procyclic forms.

Depletion of PUF3 in Lister 427 strain bloodstream-form trypanosomes had only a marginal effect on growth (Fig S3 A, B) and gene deletion was also possible (Fig S3 C, D). Induction of RNAi in pleomorphic EATRO1125 bloodstream-form trypanosomes (EATRO1125 strain) also resulted in extremely mild growth retardation (Fig 3A), but once again, clones lacking both genes were readily obtained (Fig 3B, Fig S3). Three independent clones lacking *PUF3* had the same growth rates as cells containing both, or only one *PUF3* gene, all having doubling times between 6.9 h and 7.5 h. This suggested that if PUF3 has any role in cultured bloodstream forms, its loss can be compensated during clonal selection.

**Figure 3.**
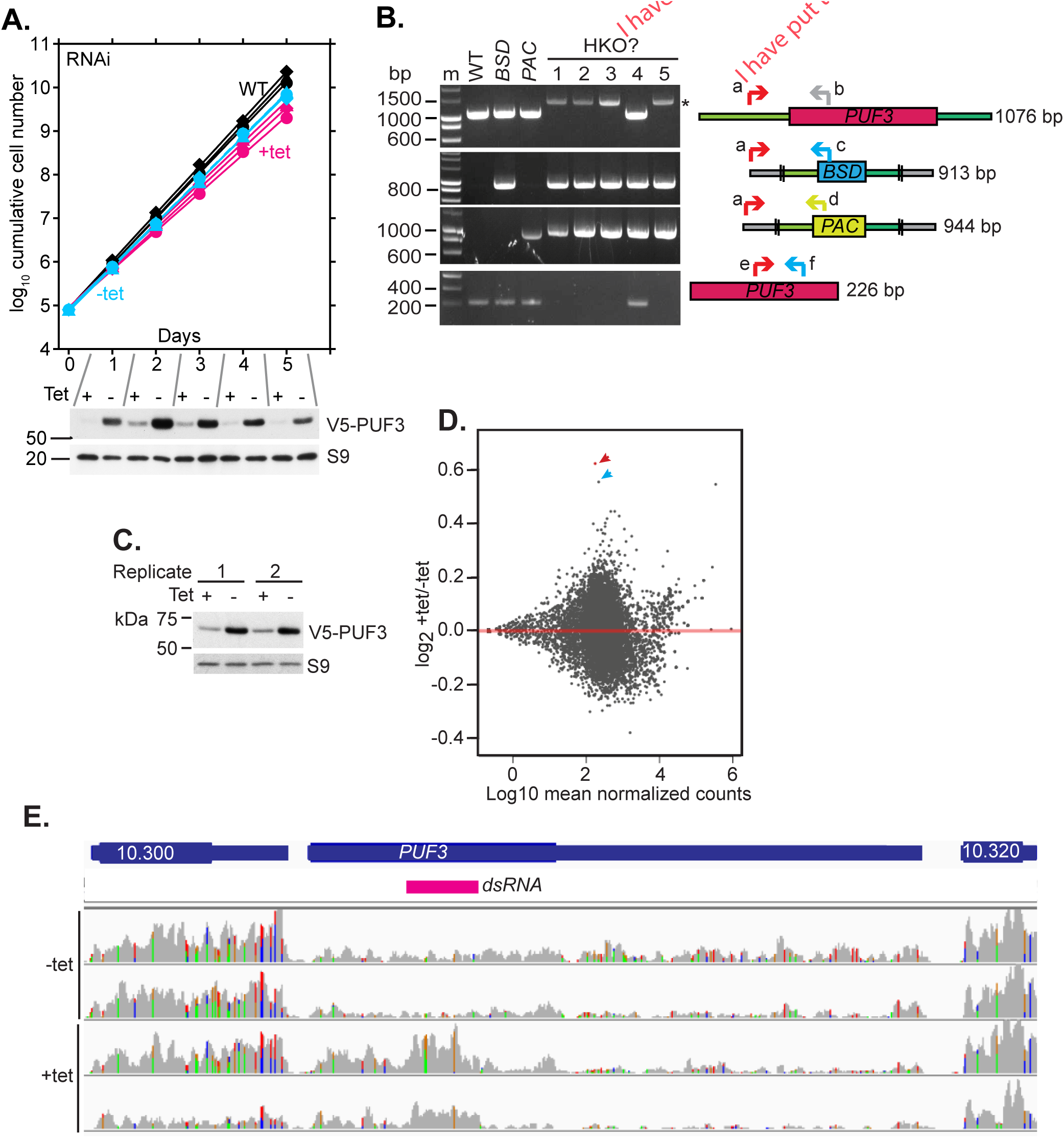
PUF3 is not essential for growth and survival of pleomorphic bloodstream forms. **A)** Bloodstream-form EATRO1125 trypanosomes with inducible RNAi targeting PUF3 were grown with or without tetracycline and diluted as necessary. Three replicates are shown. “WT” is the parental line without RNAi. A Western blot showing depletion of V5-PUF3 from one of the replicates is shown below, with ribosomal protein S9 as loading control. The remaining replicates are shown in S3 Fig. **B)** PCR test for *PUF3* knockout. Genomic DNA from wild-type, single knockouts (*BSD* and *PAC*) and candidate homozygous knockout (HKO) clones were used in a polymerase chain reaction. Double stroke lines represent the ends of the cloning fragment used for transfections. Oligonucleotides were: a) CZ6220; b) CZ3542; c) CZ5153; d) CZ5155; e) CZ6328; f) CZ6329 (S4 Table). PCR fragment sizes are on the right. Clone 4 did not have a homozygous deletion. For primers “a” and “b”, the identity of the slowly-migrating band (asterisk) is unknown, but gene deletion was confirmed by the absence of bands for primers “e” and “f” **C)** Western blot to detect PUF3 after 24h RNA interference induction, for the samples sent for RNASeq. Ribosomal protein S9 served as a loading control. **D)** RNA reads that aligned to coding sequences were analysed by DEseq2 and fold changes (+/- tetracycline) represented on a scatter plot relative to the total number of normalized counts. The red arrow points to the only significantly increased mRNA; the cyan arrow is *PUF3*. **E)** Distribution of reads mapping on the regopn of the genome encoding PUF3. Bam files generated from *PUF3* RNAi reads were loaded in Integrative Genomics Viewer (IGV version 2.4.14) using default settings. The coverages were viewed using the same data range for all the bam files and screenshots taken. The cartoons in blue are the gene models of *PUF3* (with untranslated regions adjusted to correspond to the RNA read coverage) and flanking genes while the magenta bar represents the PUF3 region targeted by RNAi.

To investigate why EATRO1125 cells with *PUF3* RNAi had slightly slower growth, we examined their transcriptomes. RNA was extracted one day after tetracycline addition. Depletion of V5-PUF3 protein was confirmed (Fig 3C), but unexpectedly, the number of reads for PUF3 was about 1.5- fold increased (Fig 3D, Table S3). In this case, therefore, the RNAi appeared to have stopped translation without complete mRNA destruction. To investigate this we examined the read distribution in the region encoding *PUF3*. Without RNAi, the sequence reads were fairly evenly distributed (Fig 3E), as expected for non-poly(A)-selected transcriptomes. In contrast, after RNAi there was an accumulation not only over the region covered by the dsRNA, but also towards the 5’- end of the transcript (Fig 3E). Comparison with the neighbouring gene (Tb927.10.300) shows that the read density in the 5’ part of *PUF3* was at least two-fold higher than in uninduced cells; it also clearly exceeded that seen 3’ to the dsRNA fragment. We have no explanation for this, but a similar discrepancy between mRNA and protein levels was seen after RNAi targeting the *CPSF30* mRNA. In that case, the *CPSF30* mRNA migrated slightly slower than without RNAi and the amount slightly increased, although the protein amount was strongly reduced [10].

Only a single additional mRNA (Tb927.9.10170, encoding anaphase-promoting complex subunit 11) was significantly increased after PUF3 depletion but the difference was only 1.5-fold (Fig 3D, Table S3). The read density over Tb927.9.10170 was uniform in all replicates (not shown), but given the total number of genes considered the increase almost certainly reflects random variation. mRNAs that were at least 2-fold enriched in the PUF3-bound fraction did not increase after RNAi (Fig S2 C). We concluded that depletion of PUF3 for 24h has no significant effect on the transcriptome.

### PUF3 may assist bloodstream-to-procyclic form differentiation

We next tested whether PUF3 has a role in differentiation of bloodstream forms to procyclic forms. Using Lister 427 trypanosomes, RNAi was induced 24 h before cells attained a density of 1.8-2.0 ×10^6^ cells/ml. Then, to induce differentiation, cis-aconitate was added and the temperature was reduced from 37°C to 27°C. Depletion of PUF3 caused a slight delay in appearance of EP procyclin (Fig S5 A). The defect was greatly accentuated in Lister 427 trypanosomes completely lacking PUF3 (Fig 4A). To check that the defect was due to to the lack of PUF3, we inducibly re-expressed the (untagged) protein. In one cloned line, re-expression of *PUF3* mRNA was tetracycline-dependent, and in the other expression was constitutive (Fig 4B). Restoration of EP procyclin expression correlated with PUF3 expression (Fig 4C). Interestingly, C-terminally myc-tagged PUF3 could not complement the defect (Fig 4D). N-terminally myc-tagged PUF3 was extremely poorly expressed (Fig S5 B) so was not tested.

**Figure 4.**
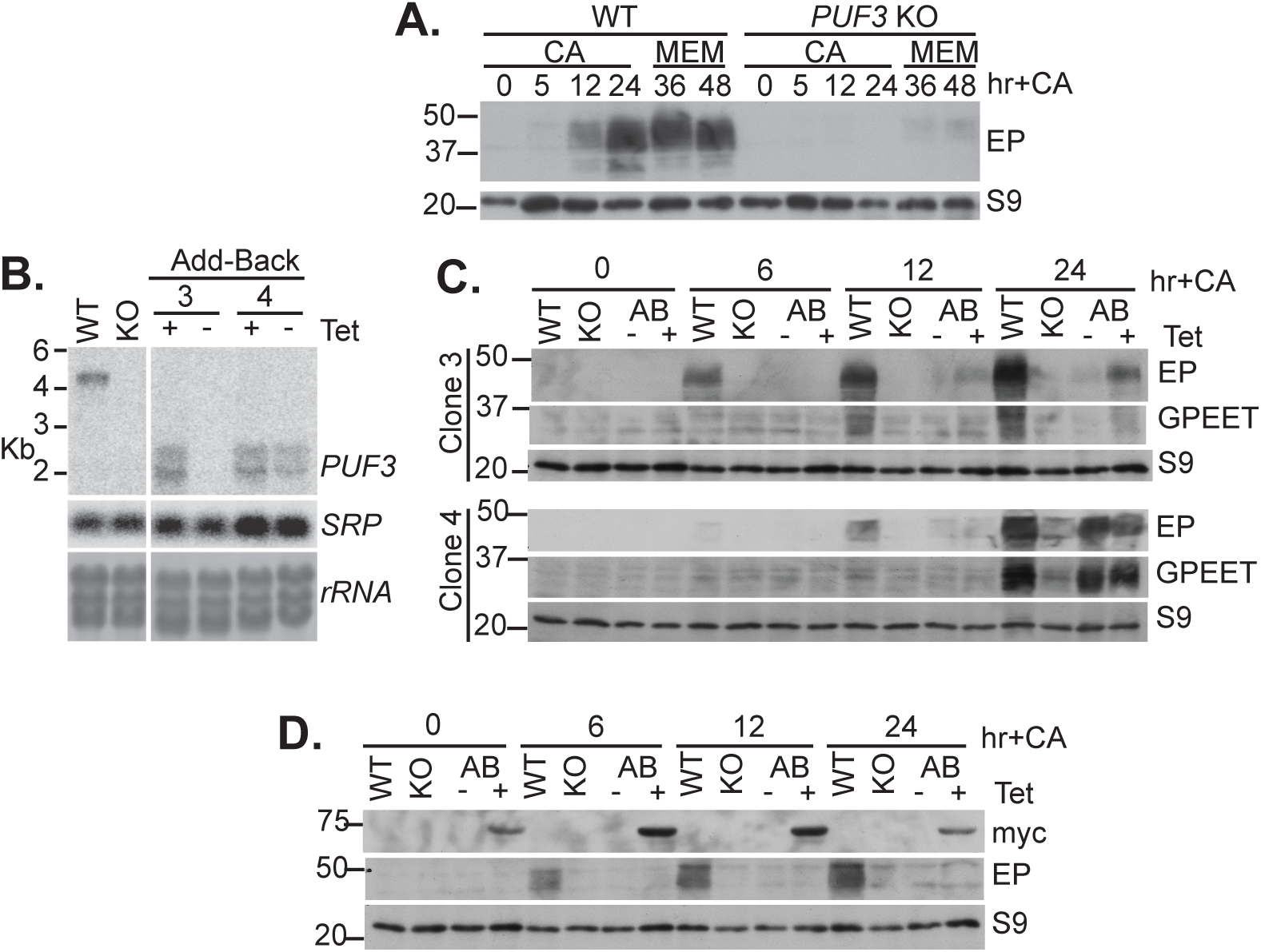
PUF3 is required for procyclin expression by differentiation-stimulated Lister 427 bloodstream forms. **A)** Monomorphic bloodstream forms were grown to 1.8-2.0 ×10^6^ cells/ml, then cis-aconitate (CA) was added to the culture (final concentration 6 mM) and the temperature was reduced from 37°C to 27°C. After 24h the medium was changed to standard procyclic medium (very-low-glucose, high-proline minimal essential medium, MEM). The time in hours after CA treatment is indicated. EP procyclin, and ribosomal protein S9, were detected by Western blotting. WT: cells with wild-type complement of *PUF3* genes; KO: cells lacking *PUF3*. **B)** Northern blot showing inducible expression of *PUF3* mRNA in the add-back clones 3 and 4. The 7SL signal recognition particle RNA (SRP) and rRNA were detected as loading controls. **C)** As in (A), but cells with inducible expression of PUF3 (“add-back”, AB) were also included, either with (+) or without (-) tetracycline addition (final concentration 100 ng/ml) 24h before the start of the experiment. The add-back results are for two different clones, clone 3 and clone 4. GPEET in addition to EP and S9 were detected by Western blotting. **D)** Expression of PUF3-myc did not complement the defect in differentiation. The experiment is similar to that in (C) except that PUF3-myc was expressed.

Next, we followed differentiation in pleomorphic EATRO1125 cells with PUF3 depletion or knock-out (Fig 5, Fig S5C). These were first grown to high density in methyl cellulose-containing media to promote stumpy-form differentiation, with or without tetracycline, and then switched to differentiation conditions (Fig 5A). Again, expression of EP procyclin was delayed after *PUF3* RNAi. In these cells, we could also see that *PUF3* RNAi caused reduced expression of the stumpy-form marker PAD1 (Fig 5B, Fig S5 C). Despite these results, however pleomorphic EATRO1125 cells that completely lacked PUF3 expressed EP procyclin normally during differentiaton (Fig 5C). Not unexpectedly, expression of PUF3-myc made no difference (Fig 5C). Preliminary results confirmed PAD1 expression in the cells without PUF3 (Fig S5 D). These results suggest that during selection of the pleomorphic knock-out clones, loss of PUF3 had been compensated for by some other unknown changes. This is not unprecedented: for example, *T. brucei* lacking UMP synthase were initially avirulent for mice, but recovered their virulence after prolonged *in vitro* culture [11].

**Figure 5.**
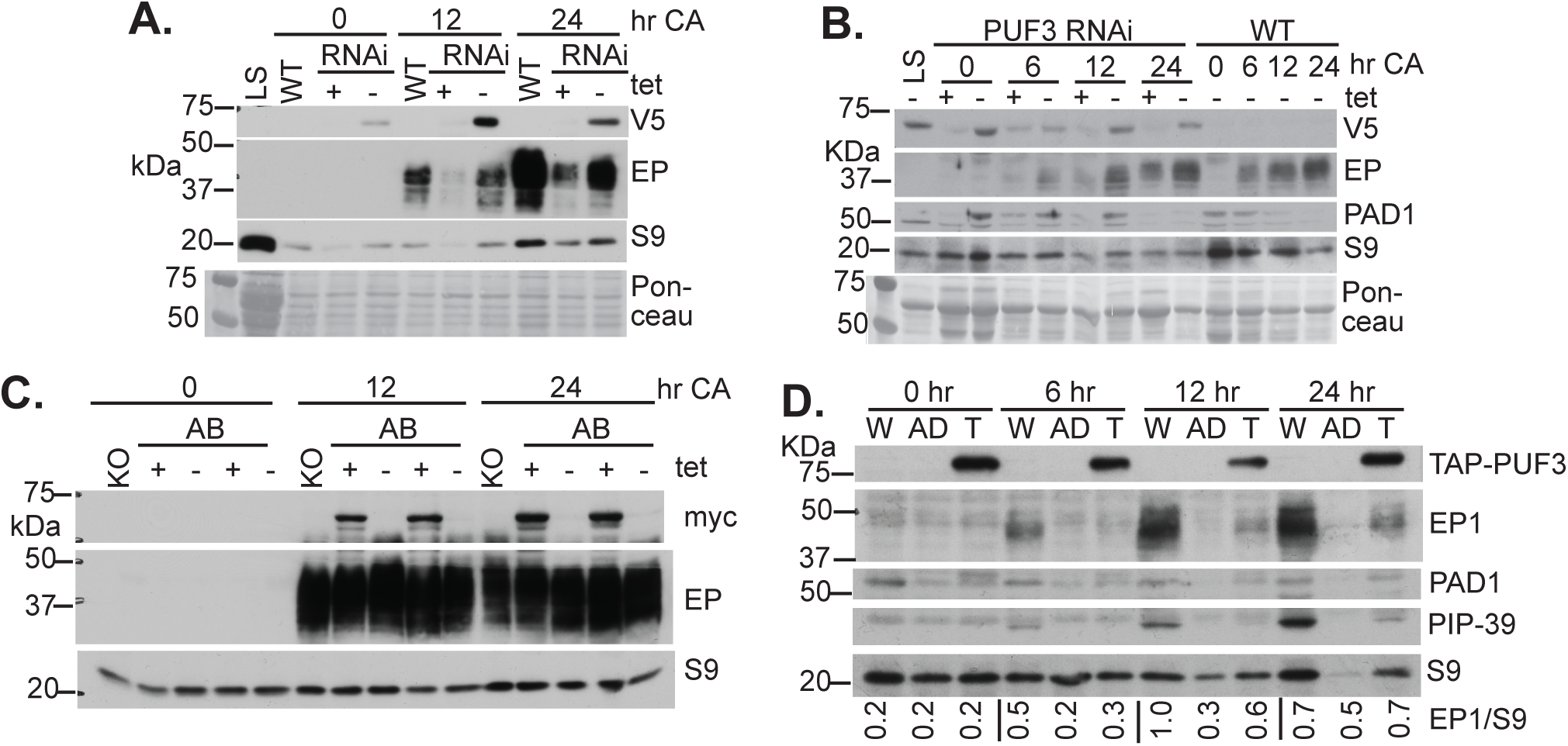
Differentiation of PUF3-depleted EATRO1125 bloodstream forms. **A)** Expression of EP procyclin during differentiation of cells with *PUF3* RNAi. RNAi was induced in EATRO1125 bloodstream forms expressing V5-PUF3, at a density of 5 x 10^5^, in methyl cellulose-containing medium. 48 hours later, cis-aconitate (CA) was added and the temperature changed to 27°C. LS: long slender cells, WT; wild-type. Ribosomal protein S9 and the Ponceau protein stain serve as controls. **B)** Replica of (A) with detection of PAD1. **C)** Cells adapted to the absence of PUF3 are not defective in EP expression in response to differentiation stimuli. The experiment is as in (A), but here cells completely lacking PUF3, with (+) or without (-) inducible expression of PUF3-myc, were used. **D)** The experiment is as in (A) but here PUF3 single knockout cells expressing *Drosophila* adenosine deaminase-PUF3 (AD) or TAP-PUF3 from the *PUF3* locus were used.

Given the potential function of PUF3 in differentiation, and the impact of tagging (Fig 4D), we checked the differentiation capability of our Lister 427 trypanosomes expressing only TAP-tagged PUF3. Another line was also included, in which all PUF3 was N-temrinally fused with *Drosophila* adenosine deaminase (AD). Cells with TAP-PUF3 expressed EP procyclin, PAD1 and the phosphatase PIP39 (Fig 5D) faster than those with AD-PUF3 but slower than wild-type. The delay could be due to the presence of lower than normal amounts of PUF3, but might also reflect an effect of the tag.

### Correct expression of PUF3 in procyclic forms is required for optimal growth

Pleomorphic bloodstream-form trypanosomes lacking PUF3 could differentiate, and grew as procyclic forms when we transferred them to appropriate medium (Fig 6A, B). Cells with inducible PUF3, but without tetracycline, grew similarly. However, if expression of PUF3 was induced during and after differentiation, growth was repressed (PUF3+ in Fig 6A, B). To examine the phenomenon in more detail, we studied the effect of expression of untagged PUF3 in well-established procyclic forms. Despite an adaptation time of several weeks, both cells lacking PUF3 and the uninduced line grew slower than wild-type (Fig 6C). However, induction of PUF3 transgene expression did not correct the defect: instead, growth was repressed (Fig 6C). This suggests that the correct level of PUF3 is required for optimal growth of procyclic forms.

**Figure 6.**
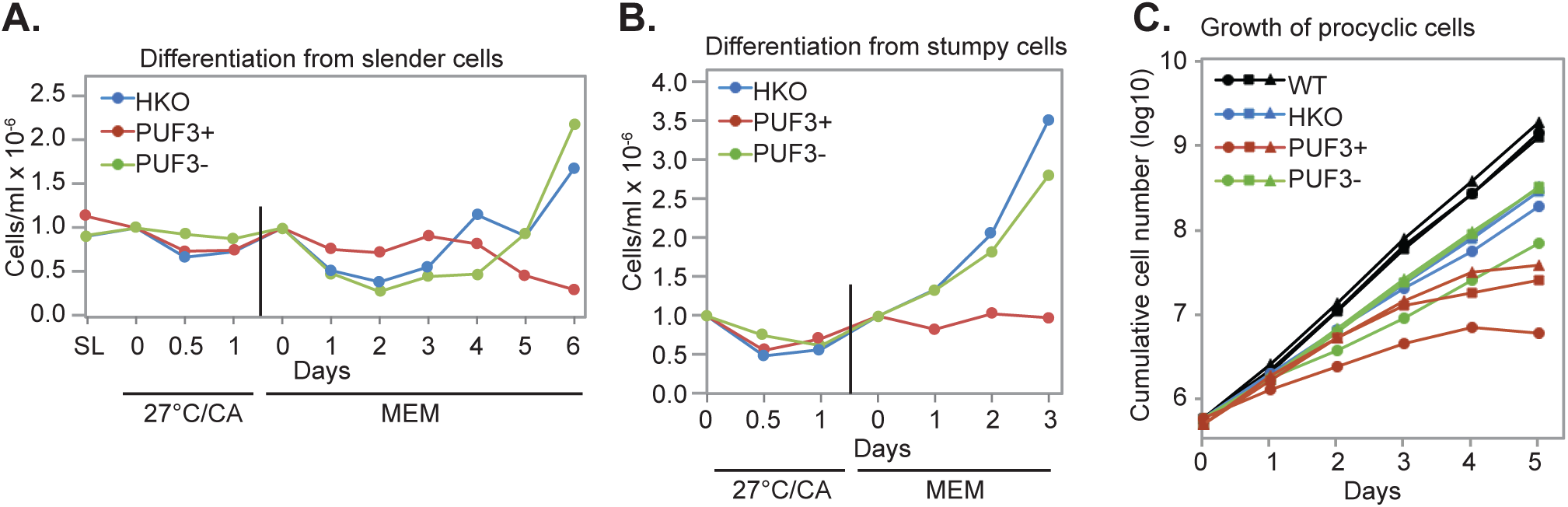
Correct expression of PUF3 is required for normal growth of procyclic forms Three different EATRO1125 cell lines were used: wild-type (WT); lacking endogenous PUF3 genes (HKO), and HKO cells with inducible expression of PUF3, grown either with (PUF3+) or without (PUF3-) tetracycline. **A)** Cells were grown to 10^6^/ml, then cis-aconitate was added, with or without tetracycline, and the temperature was reduced to 27°C (27°C/CA). After 24h the cells were transferred to procyclic-form medium (MEM). **B)** As (A) except that cells were kept at high density to induce differentiation to stumpy forms before differentiation to procyclic forms was induced. **C)** Differentiated cells were grown as procyclic forms for two weeks, then tetracycline was added to induce PUF3 over-expression. Results are for three independent cultures, indicated by the different symbols. Differentiated bloodstream cells were grown for at least four weeks as procyclics before tetracycline addition to induce PUF3 expression.

Finally we investigated whether PUF3 is required for differentiation of procyclic forms to bloodstream forms. This can be tested *in vitro* only by ectopic expression of the RNA-binding proteins RBP6 [12] or RBP10 [7]. RBP10 expression in procyclic forms forces them to differentiate to metacyclic-like forms without intervening stages and very small proportion of these metacyclic forms can replicate when placed in bloodstream-form culture conditions [7]. We therefore transfected bloodstream-form EATRO1125 cells lacking *PUF3* genes (Fig 3B) with a construct for inducible expression of RBP10-myc [7]. Regulated expression was tested, then the cells were differentiated to procyclic forms. One clone survived. The cells were cultured for 2 weeks, then RBP10-myc expression was induced with tetracycline. Two days later the cells were placed in bloodstream-form culture conditions. After 2 weeks, replicating *PUF3* HKO bloodstream-form trypanosomes were readily detectable, indicting that procyclic forms lacking PUF3 are capable of RBP10-induced differentiation to the bloodstream form.

## Discussion

Our results suggest that PUF3 is not essential for survival of cultured bloodstream-form or procyclic-form trypanosomes. Depletion of PUF3 resulted in defects in bloodstream-form growth and differentiation, but after the more prolonged culture that was required to select cells completely lacking *PUF3* genes, the cells behaved normallyy. We suggest, therefore, that the functions of PUF3 can be taken over by changes in the levels of, altered modification of, another protein - most likely, another member of the PUF family. This would not be unusual - the functions of Puf proteins also overlap in *Saccharomyces cerevisiae* [13]. In *T. brucei*, the most closely related proteins to PUF3 are PUF1 and PUF4 (S1 Fig, Fig 7). RNAi targeting pairs of these three mRNAs, however, had no effect on bloodstream-form trypanosome growth - although reduction in the proteins was not assessed [14]. PUF1 also suppresses expression when tethered, and was non-essential in procyclic forms, but pull-down and microarray results suggested that it binds to retroposon sequences [14]. It is probably much more abundant than PUF3 [15]. PUF5, like PUF3, is non-essential in procyclic forms [16], but PUF5 stimulated when tethered [4]. Tethering of PUF4 fragments yielded no evidence that it has suppressive function [4], but several other PUFs functioned only at full length [3]. The only other cytoplasmic PUF protein that is known to have suppressive function is PUF2, which is essential in bloodstream forms [17]; its binding specificity is unknown. PUF2 probably has an abundance similar to that of PUF3 [15] so a period of adaptation might be required if it were to compensate for the absence of PUF3. An alternative explanation is that if PUF3 antagonises the action of another protein, levels of that protein may have decreased after PUF3 loss.

**Figure 7.**
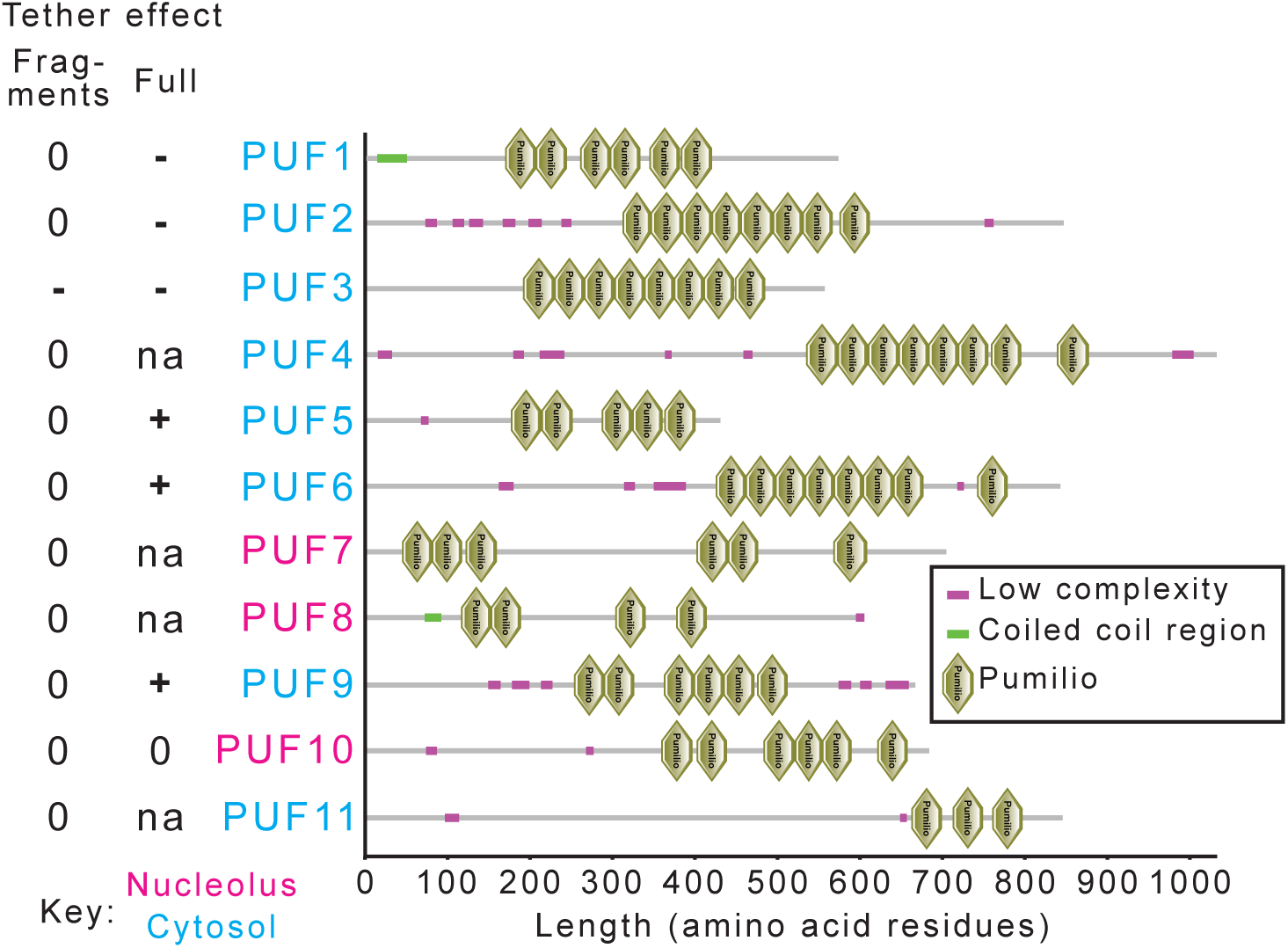
Domains of *T. brucei* PUF proteins. Key to symbols and colours are on the Figure. The effects of tethering to a reporter mRNA are shown on the left. “Fragments” refers to results from a high-throughput screen using random genomic fragments [4], and “full” refers to a screen using full-length open reading frames [3]; “na” indicates that the open reading frame was not sufficiently represented. The colours of the labels refer to the location, usually for GFP fusion proteins [2].

Ectopic expression of PUF3 had no effect in bloodstream forms but was deleterious in procyclic forms. Since we have no antibody to PUF3, we cannot judge the extent of over-expression in either form. The effect in procyclic forms is difficult to interpret. Binding of proteins to mRNAs depends on both the binding affinities and concentrations, so over-expression of PUF3 may cause it to bind to mRNAs that have sub-optimal recognition sequences. This might then displace other PUF proteins from their target mRNAs and thus disrupt regulation. Alternatively, PUF3 may be needed in order to fine-tune expression of some mRNAs in procyclic forms, in which case excess PUF3 could cause excessive repression of genuine targets.

PUF3 can suppress expression of an mRNA when tethered. Results from the previous tethering screen suggested that the region of PUF3 that is responsible for this is towards the N-terminus, with the strongest and most reproducible suppression being from fragments that started just after the initiation codon. Two short N-terminal sequences are unique to PUF3 and conserved throughout Kinetoplastea: they are WTV[QH][QE]D[DE][TYC] at residues 3-10, and YT[KQ][SC]SQVF at residues 38-45. It is possible that one or both sequences are implicated in suppression. The mechanism, however, remains unknown since no protein partners were identified. Our affinity purifications for RNASeq and mass spectrometry were done using N-terminally TAP-tagged protein from bloodstream forms. This tagged version only partially complemented the differentiation defect in cells lacking wild-type PUF3, so it is possible that the wild-type N-terminus is required for protein-protein interactions. Disruption of RNA binding seems less likely since the puf domains are separated from the N-terminus by 200 residues - but interference in the folded protein is still possible. We found no evidence that PUF3 depletion affected the transcriptome, although the cells grew slightly slower than normal; perhaps only translation is affected.

PUF3 is conserved throughout Kinetoplastea, including the free-living *Bodo saltans*, and in parasites restricted to arthropods. We therefore suggest that although we found only minor effects of depleting PUF3 in cultured trypanosomes, it may confer a selective advantage within vertebrate or invertebrate hosts.

## Materials and Methods

### Trypanosome culture and transfection

All the cells used in this study were based on stable cell lines constitutively expressing a tetracycline repressor [18], which for simplicity are referred to as wild-type. Lister 427 bloodstream forms (monomorphic) were cultured at 37°C and 5% CO_2_ in adapted HMI9 [19]: MEM including 3.24 g/L sodium bicarbonate, 23.2 mg/L bathocuprone sulphonate, 0.14 g/L hypoxanthine, 0.1 g/L sodium pyruvate, 39 mg/L thymidine, supplemented to give 1.5 mM L-cysteine and 0.2 mM β-mercaptoethanol, and with 10% fetal bovine serum (Heat inactivated at 55°C for 1 hr). EATRO 1125 bloodstream forms (pleomorphic) were cultured in HMI-9 media containing 1.1 % methyl cellulose (Sigma, M0512) to maintain their pleomorphism [20]. Densities were kept below 5x 10^6^/ml for Lister 427 and below 1×10^6^/ml for EATRO1125.

Procyclic forms were cultured at 27°C in MEM-Pros medium [21] supplemented with 5% Penicillin/Streptomycin (Gibco), 7.5 mg/l hemin and 10% of heat inactivated fetal bovine serum at a density between 10^5^ - 4×10^6^ cells/ml.

Differentiation of bloodstream slender forms to stumpy forms was done by allowing a starting density of 5-7×10^5^ cells/ml to peak at 3-4×10^6^ cells/ml, and later descend to 10^6^ cells/ml. This takes 36-48 hrs. Differentiation to procyclic forms was done by adding 6mM cis-aconitate to slender or stumpy forms at 10^6^ cells/ml and incubating at 27°C for 24 hrs before transferring the cells to MEM-Pros. Growth curves were performed using media without selecting drugs, with 100ng/µl of tetracycline when needed for transgene induction. Plasmids and oligonucleotides are listed in Table S4 and illustrated in Fig S6. Transfection of linearised plasmids and drug selections were done as described in [18].

### Immunofluorescence microscopy

Cells were induced for RNAi for 24 hrs and processed for immunofluorescence microscopy. Anti-V5 antibody (1:500, mouse) and secondary antibody with Cy3 fluorophore (1:700, anti-mouse) were used. DNA was stained with 4′,6-diamidino-2-phenylindole (DAPI). Cells were fixed in 4% paraformaldehyde at room temperature for 20 min, washed and smeared on slides. The slides were either stored at 4°C or analysed immediately. All the following steps were done at room temperature. The slides were placed in PBS for 5 min to rehydrate the cells then permeabilised using 0.2% Triton-X for 20 min. Cells were blocked with 0.5% gelatin for one hour and incubated with a primary antibody (dissolved in 0.5% gelatin) for one hour. The slides were washed three times with PBS and incubated in the dark with a secondary antibody containing a fluorescence dye for one hour. The slides were washed three times and incubated for 15 min with 1µg/ml DAPI solution. Three more washes were made and the slides air-dried before mounting with 90% glycerol-PBS. The cover slips were sealed to the slides using nail polish. Images were acquired using an Olympus IX81 microscope and analysed with Olympus xcellence software or image J.

### RNA and DNA preparation and blotting

To obtain at least 10µg of RNA, 3×10^7^ of log phase growing bloodstream-form cells (1 x 10^6^/ml for Lister 427, no more than 8x 10^5^/ml for EATRO1125) collected at 900×g for 10 min and RNA extracted using TriFast reagent according to the manufacturer’s protocol. RNA concentrations were determined using a Nanodrop spectrophotometer and the RNA stored at −80°C or used immediately. rRNAs were depleted from whole RNA extracts before RNAseq using a cocktail of DNA oligonucleotides targeting rRNAs with RNaseH [6]. For blotting, RNA was electrophoresed on a formaldehyde agarose gel, blotted on a nylon membrane by downward capillary for at least 4 hrs followed by UV cross-linking of RNA to the membrane using Stratagene® UV cross-linker. Prior to probing, the blot was blocked using pre-hybridization buffer (5×SSC, 0.5% SDS, 5× Denhardt’s solution, 100µg/ml Salmon sperm DNA). For spliced leader probing, the pre-hybridization buffer was similar to the previous one except for 6×SSC and 0.05% sodium pyrophosphate. Probes made from PCR products were prepared using PrimeIt kit (stratagene), and those from oligonucleotides using phosphonucleotide kinase (NEB) to incorporate radioactive nucleotides into the probe. The northern blot was incubated overnight with at least 2×10^6^ cpm/ml of the probe at 42°C or 65°C for an oligonucleotide or a PCR-product probe respectively. The blot was washed with buffer (2 x SSC, 0.1 % SDS) twice for 10 min per wash at room temperature and once for 15 min at 65°C then exposed to a phosphorimaging film before developing the autoradiographs with a phosphorimager (Fujifilm FLA-3000).

For Southern blotting, At least 5µg pf genomic DNA was digested using appropriate restriction enzymes and electrophoresed on an ethidium bromide-containing agarose gel. The gel was washed in hydrolysis buffer (0.25 M HCl) for 15 min, in denaturing buffer (1.5 M NaCl, 0.4M NaOH) for 15 min to denature long DNA and in neutralizing buffer (1.5 M NaCl, 0.5 M Tris-Cl pH7.5) for 15 min before blotting onto a nylon membrane and processing as described for the northern blotting.

### Chloramphenicol acetyltransferase assay

2×10^7^ cells were collected at 900×g for 10 min and washed three times with PBS. The pellet was re-suspended in 100µl of CAT buffer (100mM Tris-HCL pH 7.8) and freeze-thawed three times using liquid nitrogen and a 37°C heating block. The supernatants were collected by centrifugation at 10000×g for 5 min and kept in ice. The protein concentrations were determined by Bradford assay (BioRad) according to the manufacturer’s protocol. For each setup, 0.5µg of protein in 50µl of CAT buffer, 10µl of radioactive butyryl CoA (^14^C), 2µl of chloramphenicol (stock: 40mg/ml), 200µl of CAT buffer and 4ml of scintillation cocktail were mixed in a Wheaton scintillation tube HDPE (neoLab #9-0149) and the incorporation of radioactive acetyl group on chloramphenicol was measured using program 7 of Beckman LS 6000IC scintillation counter.

### Tandem affinity purification and mass spectrometry

6-7×10^9^ cells were collected at 900×g at 4°C for 15 min in 500 ml centrifuge containers. The cell pellet was washed twice in ice cold PBS, snap frozen in liquid nitrogen and stored at −80°C or used immediately. The cells were lysed in IPP10 buffer (10mM Tris-HCL (pH 7.8), 10mM NaCl, 0.1% IGEPAL, 50µg/ml leupeptin) by passing them through a 21-gauge needle 15 – 20 times. The lysate was cleared by centrifugation at 10,000×g for 10 min and the concentration of NaCl adjusted to 150mM before loading onto a 0.8×4 cm Poly-Prep column (BioRad) containing IgG sepharose beads that had been prewashed in IPP150 buffer (10mM Tris-HCL (pH 7.8), 150mM NaCl, 0.1% IGEPAL). This was incubated at 4°C for 2 hours with gentle rocking. The beads were washed twice with 10ml IPP150 buffer and once with 10ml TEV cleavage buffer (IPP150 buffer, 1mM DTT, 0.5mM EDTA). The beads were re-suspended in 800µl TEV cleavage buffer and incubated with TEV protease at 15-20°C for 2 hours with gentle rocking. The eluate was collected and transferred to a second column containing calmodulin beads in calmodulin binding buffer (IPP150 buffer, 10 mM β-mercaptoethanol, 1 mM MgAc_2_, 1 mM imidazole, 2 mM CaCl_2_), and incubated for 2 hours at 4°C. The beads were washed three times with calmodulin binding buffer and the proteins were eluted using 2mM EGTA. The proteins in the eluate were concentrated using trichloroacetic acid and dissolved in Laemmli buffer. This was electrophoresed for a short time so that the sample dye ran approximately 1.5cm. The gel was then submitted to the sequencing facility at ZMBH (Heidelberg) for mass spectrometry.

### RNA pull-down and RNASeq

This procedure was done in a similar manner as the TAP purification with the exception that the cells were UV-irradiated before lysis and only one purification step was done. Approximately 3×10^9^ cells were collected at 900 ×g at 4°C for 15 min, re-suspended on a petri dish in 30 ml of the supernatant on ice and UV-irradiated twice using a Stratagene UV crosslinker with an energy of 3 kilo joules. The cells were pelleted again in 4°C at 2000×g for 5 min and washed twice with ice cold PBS. Cells were lysed in IPP10 buffer consisting of KCl in place of NaCl and containing RNase inhibitor (RNasin, Invitrogen), by passing them through a 21-gauge needle 15-20 times. The lysate was cleared by centrifugation at 10,000×g for 10 min, KCl concentration adjusted to 150 mM and loaded onto a column containing prewashed sepharose beads similar as in the TAP purification. A small volume of the lysate equivalent to 7.5×10^7^ cells was collected as the input fraction and treated with 0.2mg/ml proteinase K, 8mM EDTA, 0.2% SDS at 45°C for 15 min before adding TriFast-FL reagent (Peqlab). The columns were incubated at 4°C for 2 hours with gentle rocking and the unbound lysate collected. This was also subjected to proteinase K treatment as described for the input fraction. Subsequently, the beads on the column were washed twice with IPP150 buffer and once with TEV cleavage buffer, re-suspended in 500µl of TEV cleavage buffer containing TEV protease and incubated at 15-20°C for 2 hours with gentle rocking. The cleaved product (bound) was treated with proteinase K and mixed with TriFast-FL reagent. The three fractions in TriFast-FL were stored at −80°C and later used for RNA extraction according to the manufacturer’s protocol. RNA concentrations were determined using a NanoDrop® or a Qubit® spectrophotometer and submitted to the Next-generation sequencing facility in Bioquant (Heidelberg) for library preparation. Sequencing was done at GeneCore EMBL (Heidelberg).

Fastq files were used as input in TrypRNAseq – an in-house pipeline for RNAseq (https://github.com/klprint/TrypRNAseq; 10.5281/zenodo.158920). Briefly, reads were taken through quality control, mapped on the *T. brucei* genome (TREU927) using Bowtie2, and the number of reads aligning to coding sequences (CDS) determined. Each sequence was allowed to align 20 times to allow for the repetitive nature of the *T. brucei* genome, but for statistical analyses, only one copy of repeated genes was considered [22]. To analyse the RNAi data, fold changes were compared between RNAi induced and un-induced samples using the DESeqUI adptation (10.5281/zenodo.165132) of DESeq2 [23].

The U1 and B1 libraries are incorrectly labelled in the sequence database because they were exchanged in the sequencing facility (see Legend to Supplementary Table S2, sheet 1). This was suspected from the library profiles, and became clear from the principal component analysis (Supplementary Figure S2) and from enrichment of *PUF3* reads in the B1 fraction. To further confirm the identities of the two transcriptomes we checked non-coding RNAs, which are expected to be in the unbound fractions. We re-aligned the data to all genes in the TREU927 genome, allowing each read to align only once. We concentrated on the *7SL* RNA, which was of a suitable size to be included in the sequencing library. *7SL* was indeed almost exclusively in the unbound fractions (Supplementary Table S1, sheet 4).

## Data availability

The sequence results are available under the following accession numbers: RNAi - E-MTAB-8062; RIP-Seq - E-MTAB-8068.

## Acknowledgements

Kevin Marucha was supported by a fellowship from the Deutsches Akademisches Austauschdienst (DAAD). The work was also supported by the Deutsche Forschungsgemeinshaft, Grant CL112/24. We thank Ute Leibfried and Claudia Helbig for technical assistance, and Keith Matthews for the antibodies to PAD1 and PIP39. We thank the ZMBH mass spectrometry facility for protein analysis and David Ibberson (BioQuant, Heidelberg) for RNASeq library construction.

## Supplementary Tables

***<u>S1 Table</u>***

RIPSeq data: mRNA association with TAP-PUF3 in bloodstream forms

***<u>S2 Table</u>***

Mass spectrometry data: lack of protein association with TAP-PUF3with other proteins

***<u>S3 Table</u>***

RNASeq data: effect of RNAi targeting PUF3

***<u>S4 Table</u>***

Plasmids and oligonucleotides

## Supplementary Figures

**S1 Figure.**
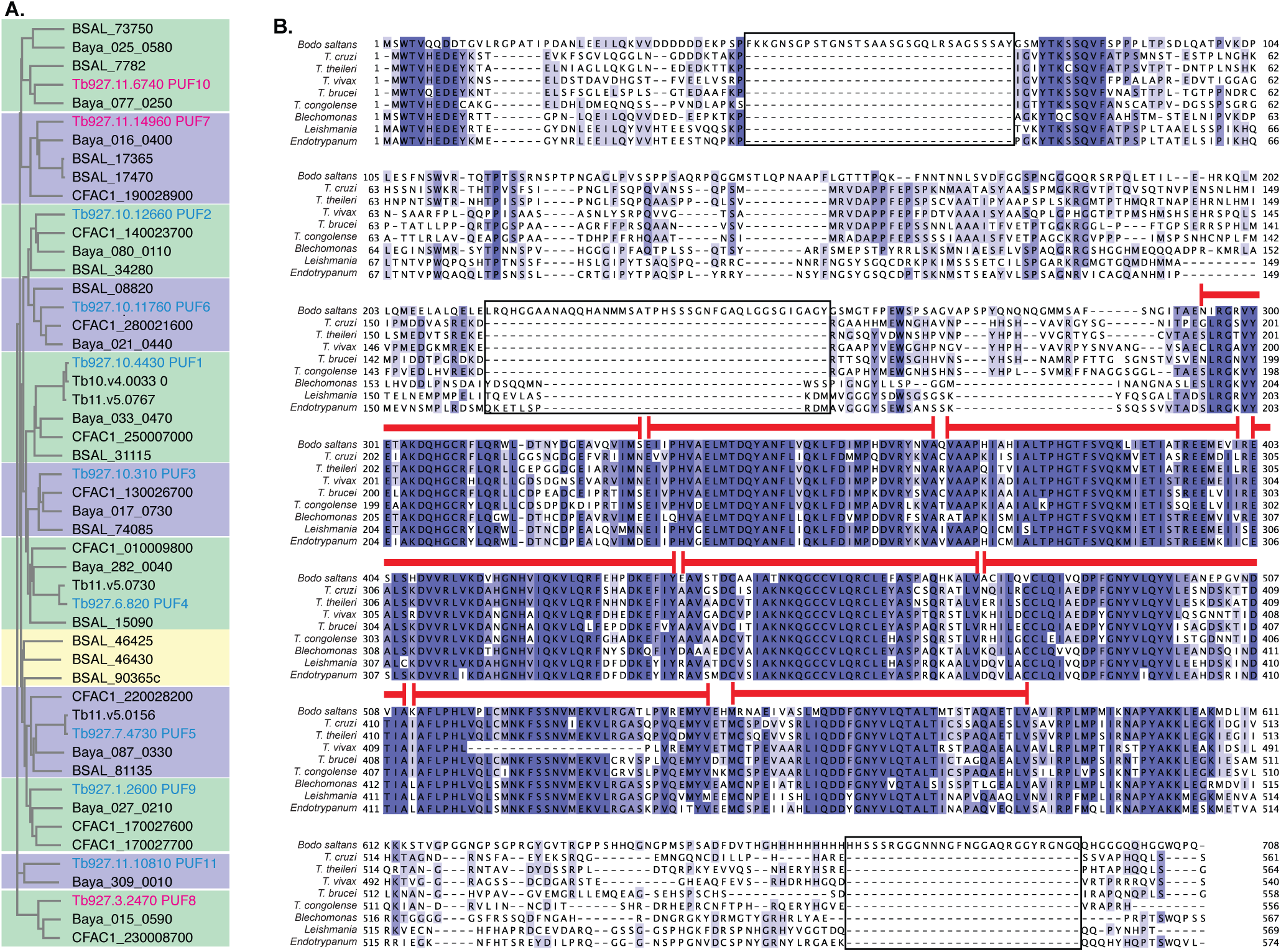
**(A)** Cladogram of PUF orthologs **(B)** PUF3 protein sequences from *Trypanosoma brucei, T. vivax, T. congolense, T. theileri, T. cruzi, Leishmania major, Blechomonas ayalai, Endotrypanum monterogeii* and *Bodo saltans* were aligned using Clustal Omega (EMBL-EBI). The shading highlights similarities. Red bars indicate the position of the PUF repeats in *T. brucei* PUF3. The apparent gap in the seventh PUF repeat for *T. vivax* is caused by a gap in the sequence assembly.

**S2 Figure.**
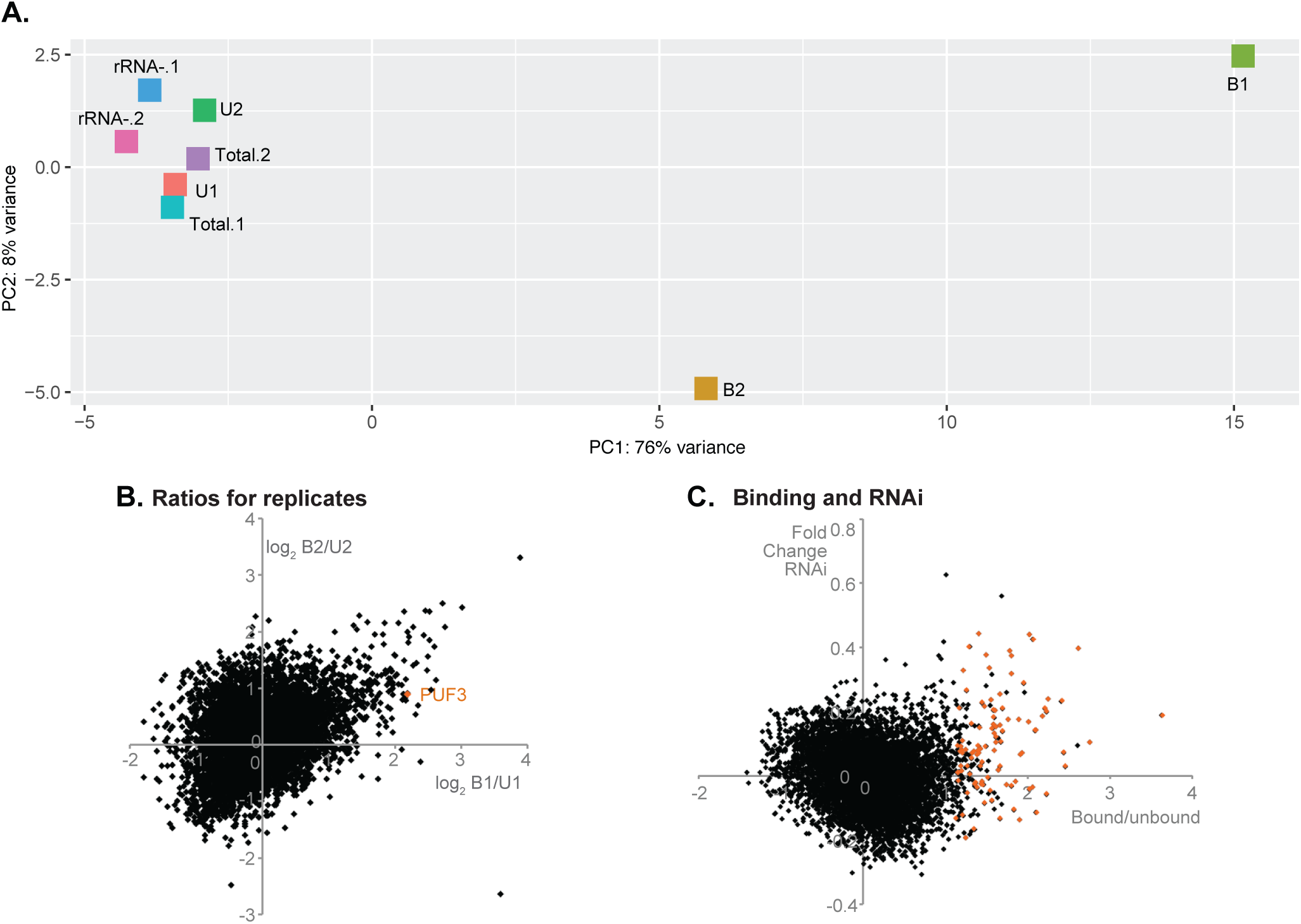
mRNA association with TAP-PUF3. TAP-PUF3 was bound to IgG beads and bound and unbound RNAs were sequenced. **A)** Principal component analysis for RNA-pull-down results. B - eluate or bound fraction; U - flow-through or unbound fraction. rRNA-is input RNA after rRNA depletion; Total is input RNA without rRNA depletion. Raw data are in Table S1. **B)** The ratios of bound/unbound reads per million were calculated for all unique genes. *PUF3* mRNA was enriched 4.6-fold in replicate 1 and 1.88-fold in replicate 2 (orange dot). There was a weak positive correlation between the two sets of enrichment values (R= 0.35 for log-transformed results). **C)** Average binding ratios for each mRNA are plotted against the fold change after RNAi. None of the RNAi effects was statistically significant. Most mRNAs with higher bound/unbound ratios showed no RNAi effect at all. Orange dots represent mRNAs for which bound/unbound was >2 in both experiments, assuming the sample swap.

**S3 Figure.**
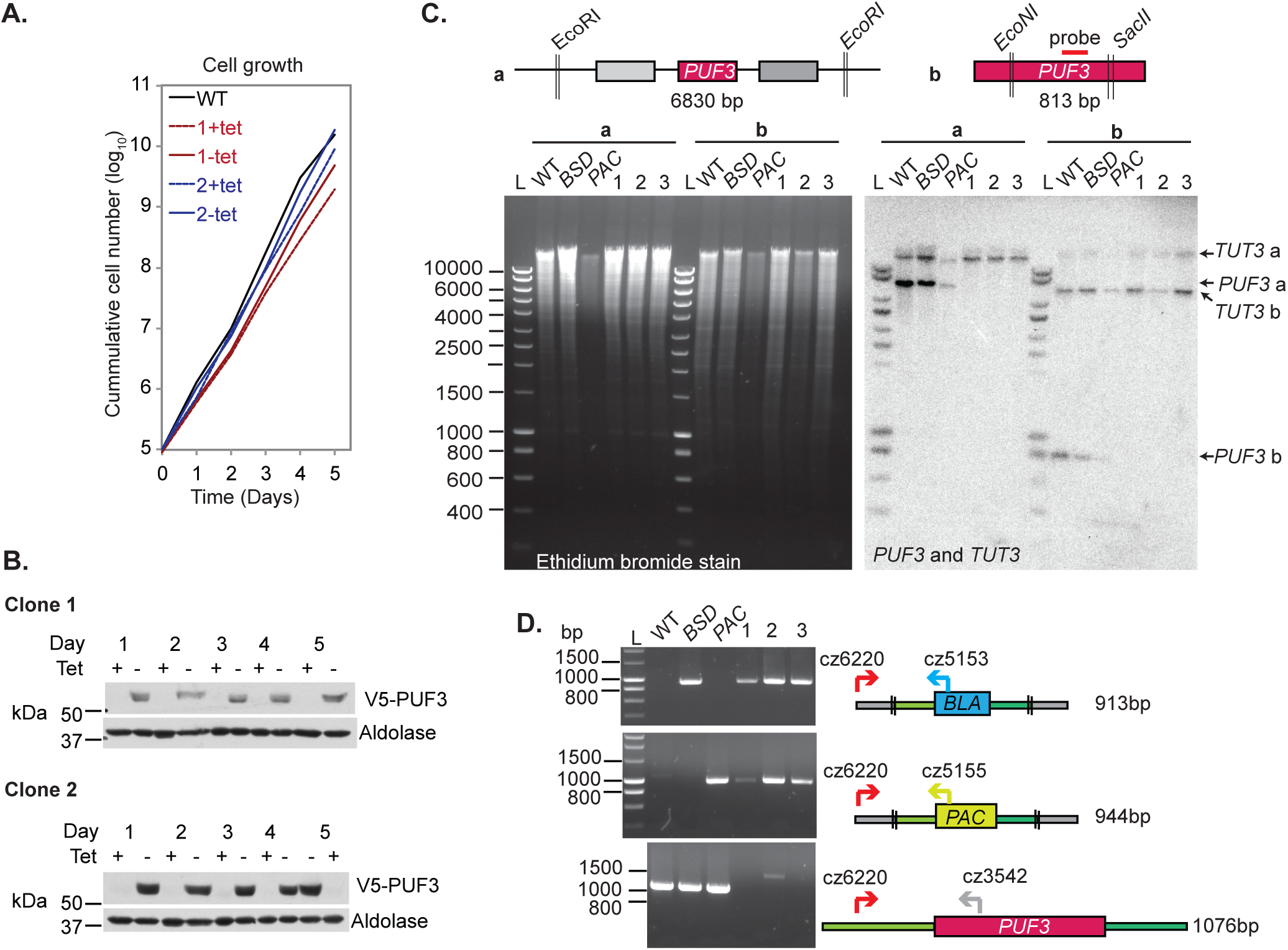
PUF3 is not essential for survival of monomorphic bloodstream forms. **(A)** Growth of bloodstream-form Lister427 trypanosomes with *in situ* N-terminally V5-tagged PUF3 and tetracycline-inducible *PUF3* RNAi. Results are shown for wild-type cells and for two independent clones grown with (+) and without (-) tetracycline. **(B)** Western blots showing the expression of a V5-tagged PUF3 in the cell lines in (A). Antibody to aldolase was used as a loading control. **(C)** The upper panel represents the design of a Southern blot. Coding sequences of genes are represented by rectangles and vertical double-strokes show restriction sites. Genomic DNA of cells selected for homozygous knockout (transfected with plasmid pHD2851 and pHD2852) was digested with *Eco* RI (experiment a) or *Eco* NI and *Sac* II (experiment b). The digested DNAs for wild-type (WT), single knockouts (*BSD* or *PAC*) and three candidate homozygous knockout clones were analysed by Southern blotting using a probe targeting *PUF3* (as indicated in the upper panel), with *TUT3* as a loading control. L: marker ladder. **(D)** DNA from (C) was used in a polymerase chain reaction using the primers shown (Table S4). Double stroke lines represent the ends of the cloned fragments used for transfections.

**S4 Figure.**
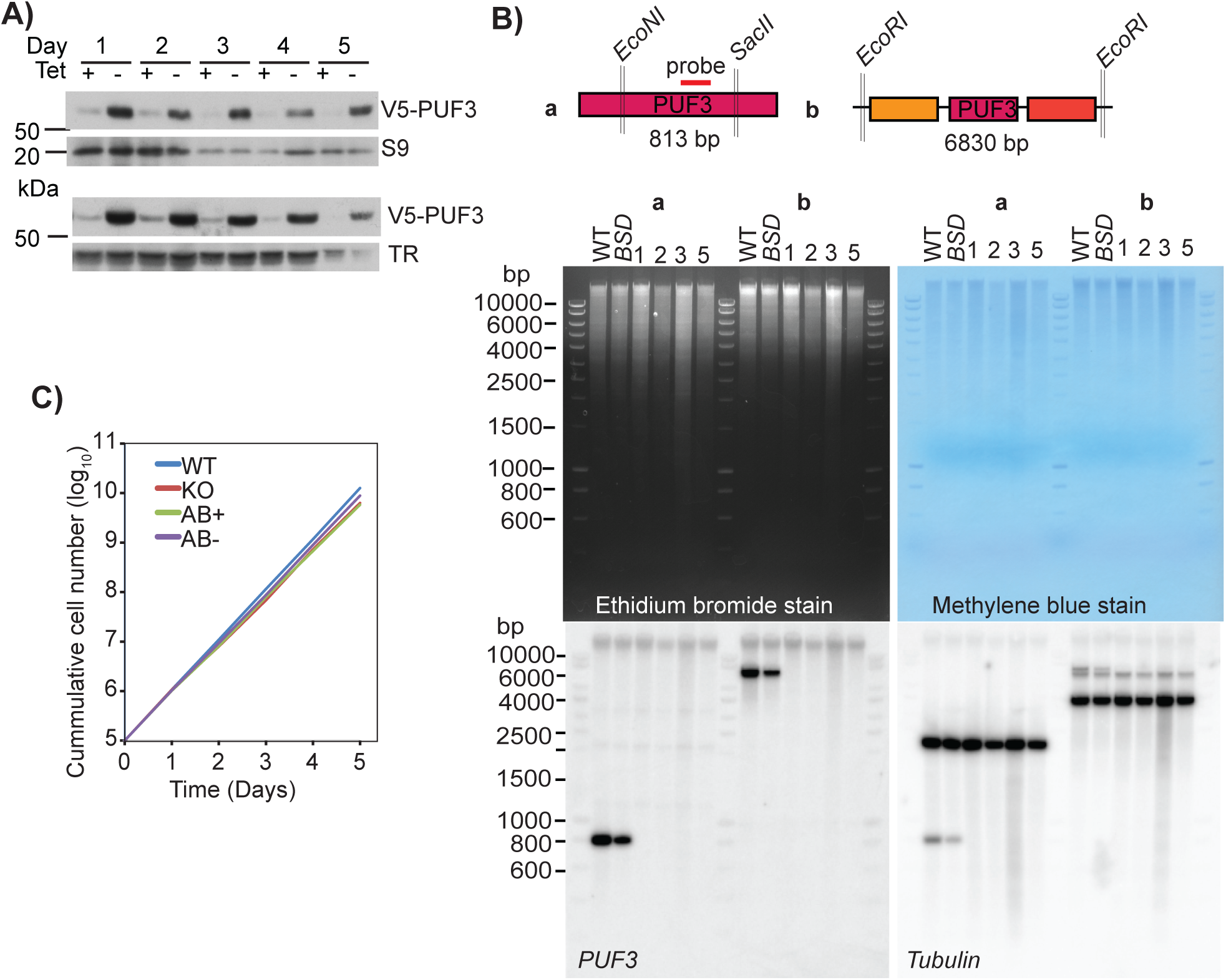
PUF3 is not essential for survival of pleomorphic EATRO1125 bloodstream forms. **(A)** Western blots showing V5-PUF depletion for two additional samples from Figure 2A. Controls are ribosomal protein S9 or trypanothione reductase (TR). **(B)** The upper panel represents the design of a Southern blot. Coding sequences of genes are represented by rectangles and vertical double-strokes show restriction sites. Genomic DNA of cells selected for homozygous knockout (transfected with plasmid pHD2851 and pHD2852) was digested with *Eco* NI and *Sac* II (experiment a) or *Eco* RI (experiment b). The digested DNAs for wild-type (WT), single knockouts (*BSD*) and four candidate homozygous knockout clones were analysed by Southern blotting first using a probe targeting *PUF3* (as indicated in the upper panel) and second with a probe for beta-tubulin. **(C)** Cumulative growth of the bloodstream forms, mean values. Variation (not shown) was so low that all of the values overlapped. AB: Add-back of tetracycline-inducible PUF3-myc, +tet or -tet.

**S5 Figure.**
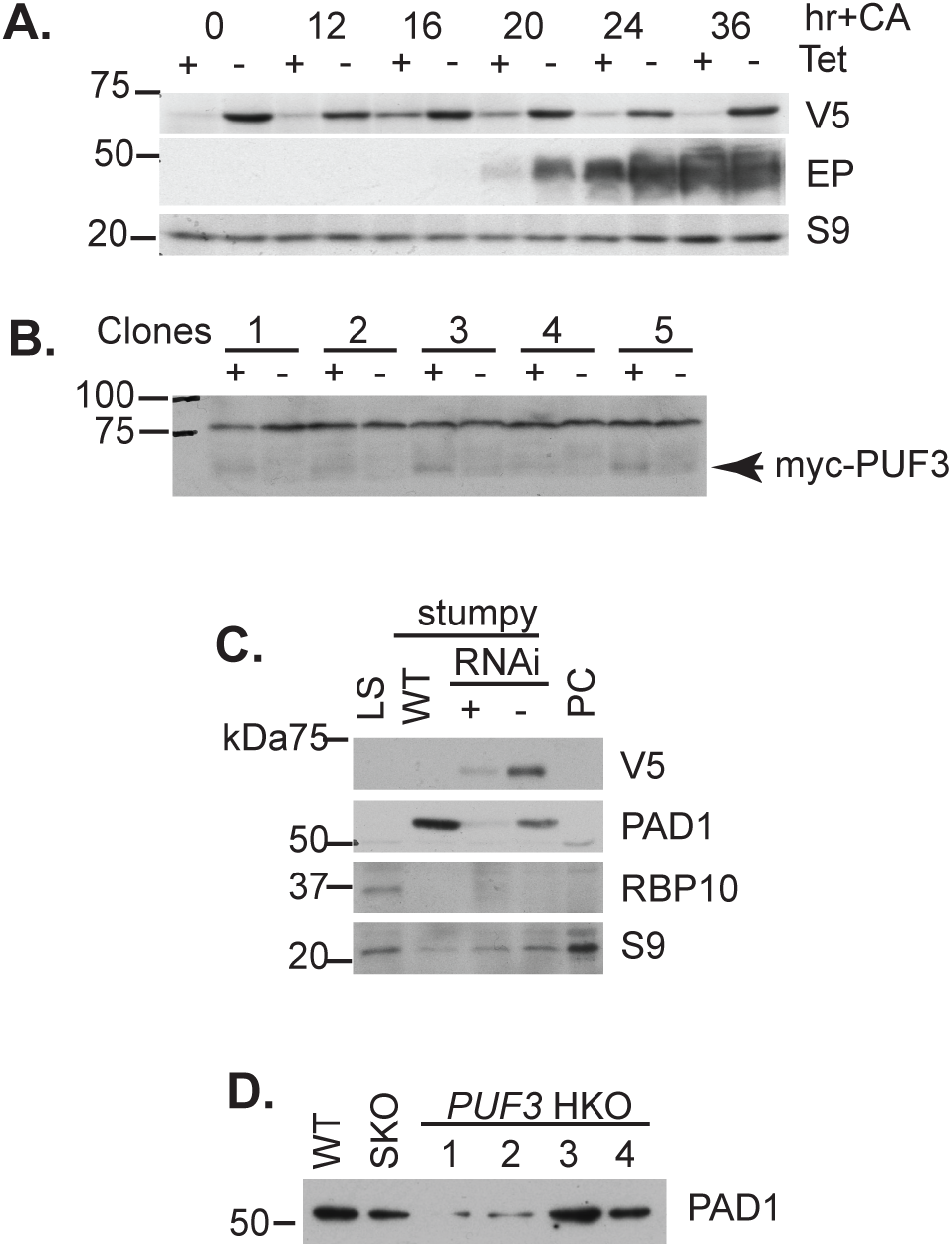
Differentiation of bloodstream forms lacking PUF3. **(A)** RNAi targeting PUF3 in Lister 427 bloodstream forms gave a transient differentiation defect. RNAi was induced 24 h before cells attained a density of 1.8-2.0 ×10^6^ cells/ml. Cis-aconitate (CA) was added to the culture (final concentration 6 mM) and the temperature was reduced from 37°C to 27°C. The time in hours after CA treatment is indicated. The Western blot shows expression of EP procyclin and V5-tagged PUF3, with ribosomal protein S9 as a control. **(B)** Inducible expression of N-terminally myc-tagged PUF3 in five Lister 427 bloodstream-form clones. The protein was barely detectable. **(C)** Expression of PAD1 and RBP10 after *PUF3* RNAi in EATRO1125 bloodstream forms. The experiment was as in Fig 5B. **(D)** Expression of PAD1 in the EATRO 1125 stumpy cells in Figure S4, which lack PUF3

**S6 Figure.**
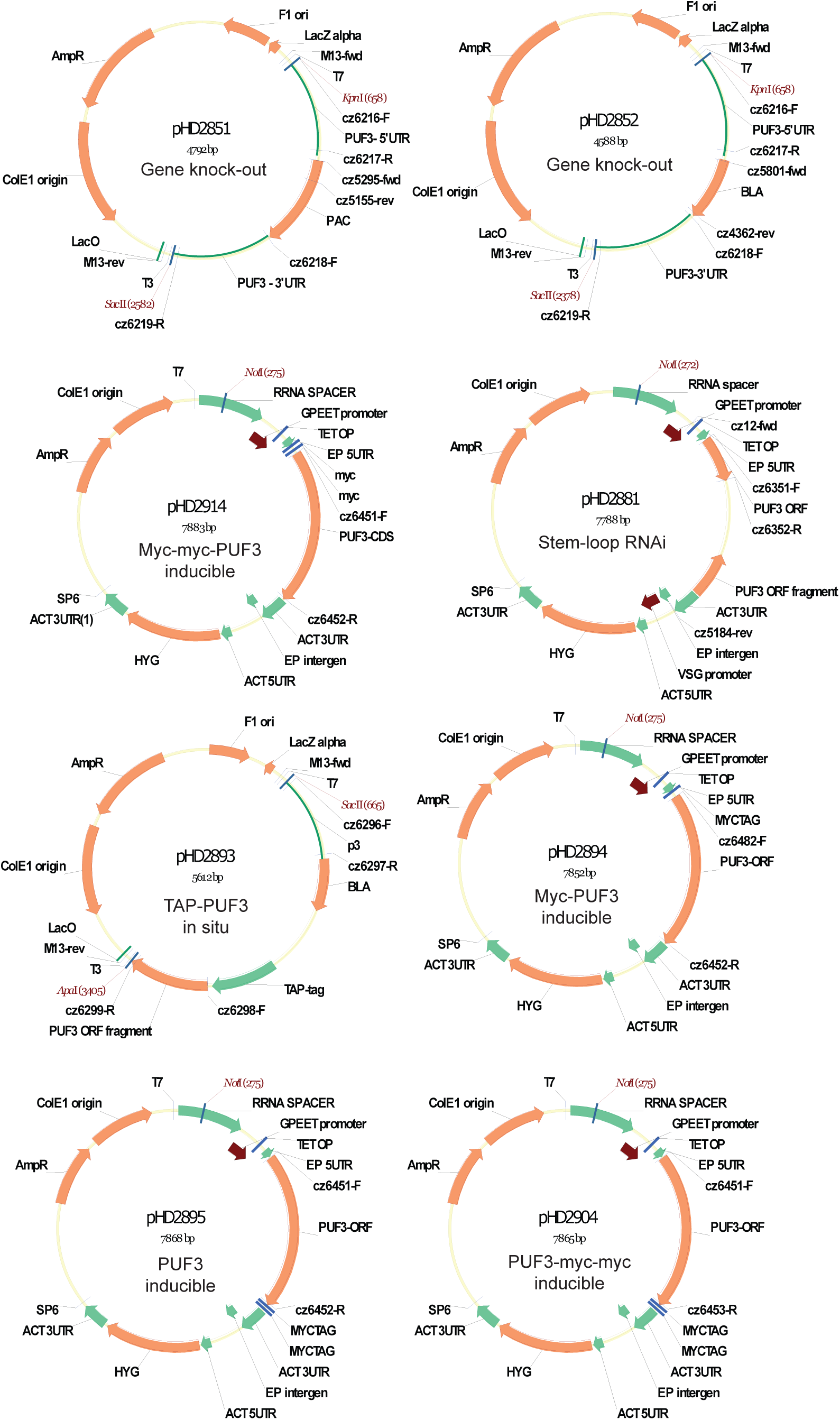
Plasmids made for this work

